# Development of Top-Down Cortical Propagations in Youth

**DOI:** 10.1101/2022.06.14.496175

**Authors:** Adam Pines, Arielle S. Keller, Bart Larsen, Maxwell Bertolero, Arian Ashourvan, Dani S. Bassett, Matthew Cieslak, Sydney Covitz, Yong Fan, Eric Feczko, Audrey Houghton, Amanda R. Rueter, Tinashe Tapera, Jacob Vogel, Sarah M. Weinstein, Russell T. Shinohara, Damien Fair, Theodore Satterthwaite

## Abstract

Hierarchical processing requires activity propagating between higher and lower-order cortical areas. However, studies of brain development have chiefly quantified fluctuations within regions over time rather than propagations occurring over space. Here, we leveraged advances in neuroimaging and computer vision to track cortical activity propagations in a large sample of youth (*n*=388). We found that propagations robustly ascend and descend the cortical hierarchy, and that top-down propagations become both more prevalent with cognitive control demands and with development in youth.

## Main

The hierarchical organization of the cortex provides a scaffold for bottom-up sensory integration and top-down control^1,2,3^. Existing evidence suggests that cortex-wide hierarchical organization is a product of protracted development^4,5,6^. Understanding the development of hierarchical processing is critical, as developmental deficits in cognitive control that are thought to rely on top-down processing are associated with transdiagnostic psychopathology^7^, reduced quality of life^8^, and youth mortality^9^. In the brain, hierarchical processing necessarily involves activity propagating through space between higher- and lower-order areas. However, most fMRI studies of hierarchical processing have chiefly quantified activity fluctuations in fixed regions over time, rather than examining activity propagations over space. While some recent work has approximated spatial propagations by detailing sequences of activations in nodes within graph-based representations of the brain^10,11,12^, non-invasive measurement of cortical activity propagations in humans has remains an open challenge, and it is unknown how propagations may evolve in development.

Several recent studies have used a combination of fMRI and intracranial recordings to demonstrate that infraslow but large-scale activity systematically propagates along a principal gradient (PG)^13^ of cortical organization from lower-to higher-order areas^14,15,16^. Two studies also noted top-down propagation, where activity instead moved from higher-to lower-order areas. Intriguingly, such top-down propagations were associated with alertness^15^ and the ascending arousal system^17^, suggesting that top-down propagations might be linked to top-down cognitive processing. However, to infer hierarchical directionality, these approaches relied upon a single, group-level cortical pattern linked to the time series of a single variable: either respiratory variability^15^, the global signal^16,18^, or the difference in signal from two subcortical regions over time^17^. While these approaches revealed prominent, stereotyped hierarchical propagations, they are circumscribed to their respective time-locked variables of interest, and predominantly reveal group-averaged propagations. As a result, little is known about how propagations vary across individuals and mature with development.

Here, we fill this critical gap by capitalizing upon a widely-used method in computer vision – optical flow – to quantify activity propagations across the cortex. Optical flow enabled us to derive *directional* information regarding propagations directly from changes in local BOLD signal (**methods**). In neuroscience, optical flow has been primarily implemented either on group-level patterns^16^, or on mesoscale sections of cortex^19^. Recently, the optical flow algorithm was adapted to efficiently estimate biological motion on the surface of spheres^20^. We leveraged this advance to quantify the movement of the BOLD signal directly on each participant’s cortex following spherical registration. We hypothesized that this approach would reveal bottom-up and top-down propagations along the principal gradient of macroscale cortical organization. Furthermore, we predicted that top-down propagations would be associated with task demands and become more prominent with age in youth. To test these hypotheses, we leveraged a large developmental dataset with both high-quality resting-state and task fMRI data^17^ (*n* = 388 after QC, mean age = 15.6, *SD* = 3.7 years).

Optical flow yielded vector fields describing the direction of signal propagation between pairs of sequential fMRI volumes mapped to the cortical surface via spherical registration (**Figure 1a**). To evaluate the presence of hierarchical propagations, we extracted the gradient vector field (∇) of an established map that defines the principal gradient (PG) of the cortical hierarchy (∇PG, **Figure 1b**). Because gradient vector fields describe the direction of image intensity increases, ∇PG describes the direction of hierarchical ascent at each point on the cortex. Local ∇PG directions were subsequently utilized as reference directions for optical flow vectors for each participant (**Figure 1c**). After removing volumes corrupted by head motion, we recorded the difference in the angle (in degrees) of the direction of activity estimated by the optical flow vectors with respect to the direction of hierarchical ascent defined by the ∇PG **(Figure 1d)**. In this framework, alignment with the angle of hierarchical ascent (0° from ∇PG) indicates a bottom-up propagation, whereas flow in the opposite direction (180° from ∇PG) indicates a top-down propagation (**Figure 1e, Figure S1, Figure S2**).

**Figure 1:**
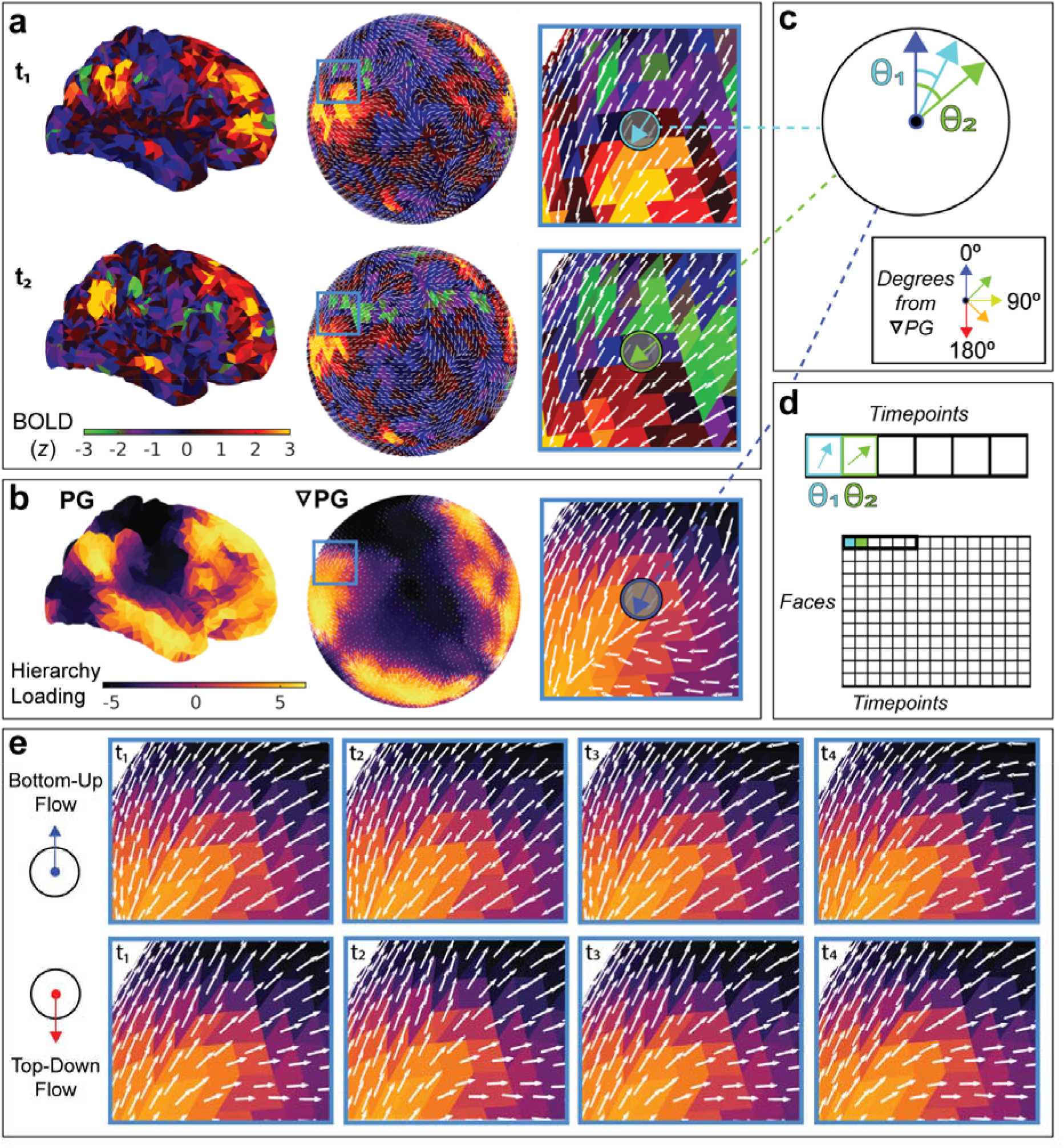
Schematic for spherical optical flow and assessment of hierarchical propagations. **a)** To estimate the spatial directionality of activity across the cortex, all fMR images are projected to the *fsaverage4* spherical surface. Specifically, for each pair of sequential images, we used optical flow to estimate the directions of signal propagations at each face on the cortical mesh. **b)** To estimate the direction of hierarchical ascent, the gradient vector field of a validated map of cortical hierarchy^10^ was extracted along the cortical surface (∇PG). This procedure yields vectors across the entire cortex, with each vector describing the most immediate direction of hierarchical ascent for any given face on the mesh. **c)** To quantify directional distributions, each optical flow direction is assessed relative to the direction of hierarchical ascent over all sequential image pairs. **d)** This procedure is repeated for each face on the cortical mesh to yield a matrix of activity directions relative to ∇PG over time for each participant. **e)** Example bottom-up and top-down propagations: vectors are extracted from pairs of sequential BOLD images (white arrows) and overlaid onto the group-level PG (yellow-black shading).

We observed a predominance of both bottom-up and top-down propagations, which formed a bimodal distribution. These bimodal distributions were evident at the group (**Figure 2a**) and participant-level (**Figure 2b**). To rigorously test whether propagations were enriched for bottom-up and top-down directionality, we used a conservative spin-based permutation method that perseveres the spatial covariance structure of the data **(Figure 2c)**. This procedure revealed that the angular distributions of propagations were specifically aligned with ∇PG for every participant in the sample, far beyond what could be expected by chance (real data median SD from null distribution = 13.6 SD; *p* < 0.001 for every participant in the sample). To further confirm that optical flow captured continuous propagations rather than differences between discrete activation patterns, we shuffled the temporal ordering of fMRI volumes from each participant **(Figure 2d)**. These temporal permutation tests confirmed that optical flow captured specific sequences of activity that were not present in shuffled data (real data median SD from null distribution = 18.8 SD; *p* < 0.01 for 93% of participants in the sample).

**Figure 2:**
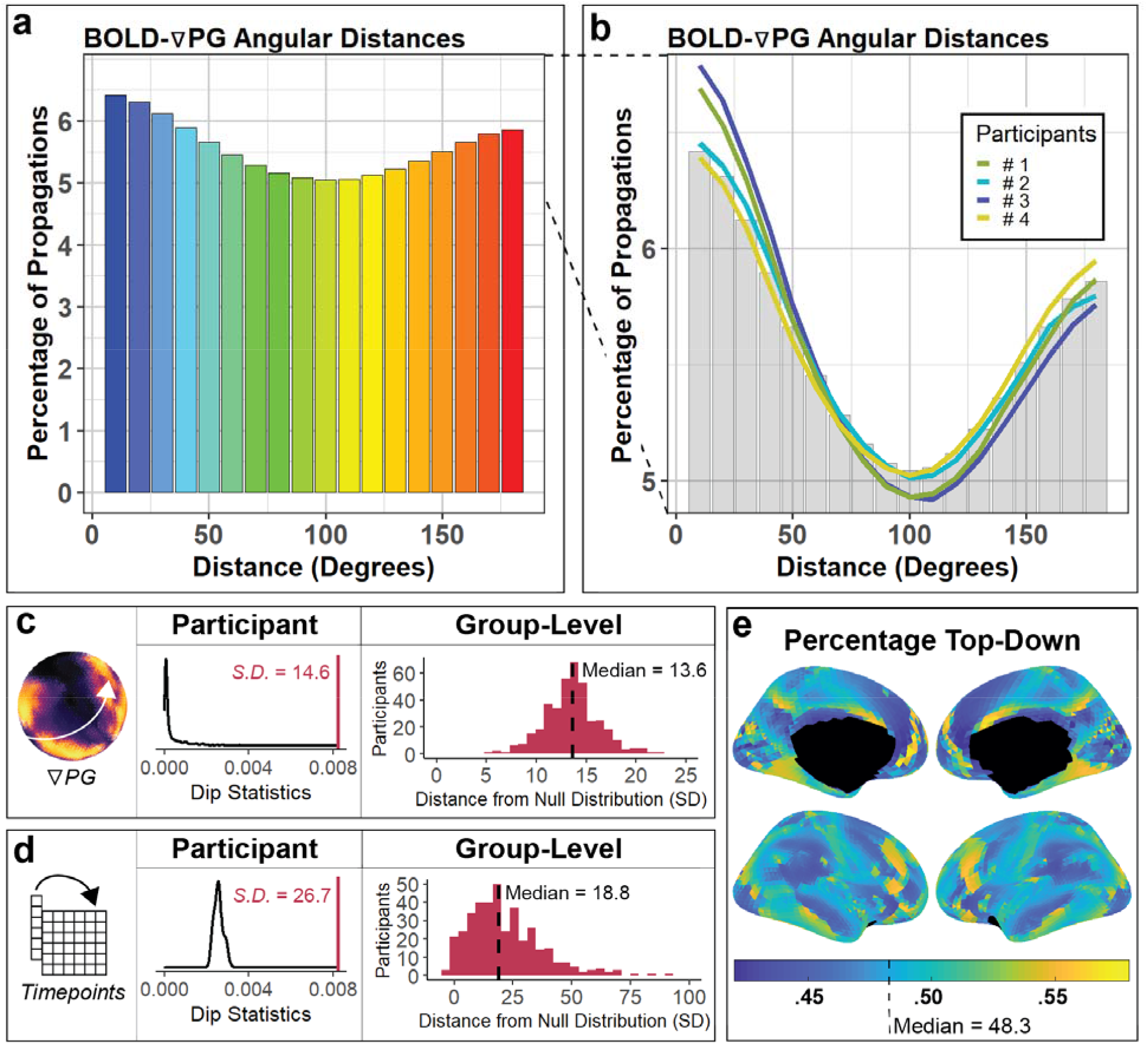
Cortical activity propagates up and down the cortical hierarchy. **a)** Group-level directional distributions revealed a bimodal distribution of angular distances between ∇PG and flow vectors (*n* = 4.4 billion optical flow directions over all TRs and participants). **b)** Directional distributions are bimodal for hierarchical ascent (0°) and descent (180°) within individual participants. The percentage axis is rescaled from panel (**a)** for detail, with the equivalent y-axis range demarcated across panels with dashed lines. **c)** Spatial null models permuted the reference directions (∇PG) continuously in space, preserving the spatial covariance structure of the original map (left). Spatial null models are computed within participants (middle; participant #1 from panel **b**) by comparing the dip statistic obtained from permuted reference directions (black distribution) and the true dip statistic (red line). Whereas 1.96 standard deviations from the mean is a common statistical threshold for significance, we found that true dip statistics tended to be approximately 13.6 standard deviations from the null distribution (right). **d)** Temporal null models involved shuffling the order of retained fMRI volumes in time, preserving complex spatial patterns found within individual images (left). Temporal null models were computed within participants (middle; participant #1 from panel **b**) by comparing the dip statistic obtained from permuted fMRI volume ordering (black distribution) and the true dip statistic (red line). True dip statistics were a median of 18.8 standard deviations from the null distribution (right). 5 participants with positive outlier values were omitted for clarity (SDs = 166.7, 189.3, 258.9, 369.7, and 747.9). **e)** All faces exhibited a mix of both bottom-up (<90°) and top-down (>90°) propagations, but certain regions were enriched for bottom-up propagations or top-down propagations. For example, bottom-up propagations were more common in medial prefrontal cortex, whereas top-down propagations were more common in dorsolateral prefrontal cortex.

Having demonstrated the presence of hierarchical propagations in all participants, we next sought to define the spatial distribution of bottom-up and top-down propagations. For each location on the cortex, we quantified the percentage of propagations that could be characterized as bottom-up or top-down (**Methods**). While all regions exhibited a mix of both bottom-up and top-down propagations at different points in time, bottom-up propagations were more common in certain regions (i.e., medial prefrontal cortex) and top-down propagations were enriched in others (i.e., dorsolateral prefrontal cortex; **Figure 2e**). At the participant-level, the percentage of top-down optical flow vectors was highly correlated with our statistical summary measure of non-unimodality (i.e., dip statistic, *r* = .70, *p* < 0.01×10^−14^; **Figure S3**). This percentage allowed us to directly test whether top-down propagations became more common under task demands and with development in youth. Specifically, we sought to evaluate whether the prevalence of top-down propagations was modulated by a cognitive task that requires top-down cognitive control. We compared propagations observed during rest to those present during a modified Go/NoGo task, where top-down control is intermittently required to suppress reflexive button-pressing^21^. Mass univariate analyses revealed more top-down propagations during task than rest (*t*_*f*ace_ = 2.37-13.97, *p*_fdr_ <0.05; **Figure 3a**). While these effects were distributed across the cortex, increases in top-down propagations were particularly prominent in regions within the dorsal and ventral attention systems. Increases were maximal in the left upper-extremity subdivision of motor cortex, likely corresponding to the uniform usage of the right hand to execute task demands across participants^21^. These results suggest that task demands modulate the prominence of top-down propagations within individuals.

**Figure 3:**
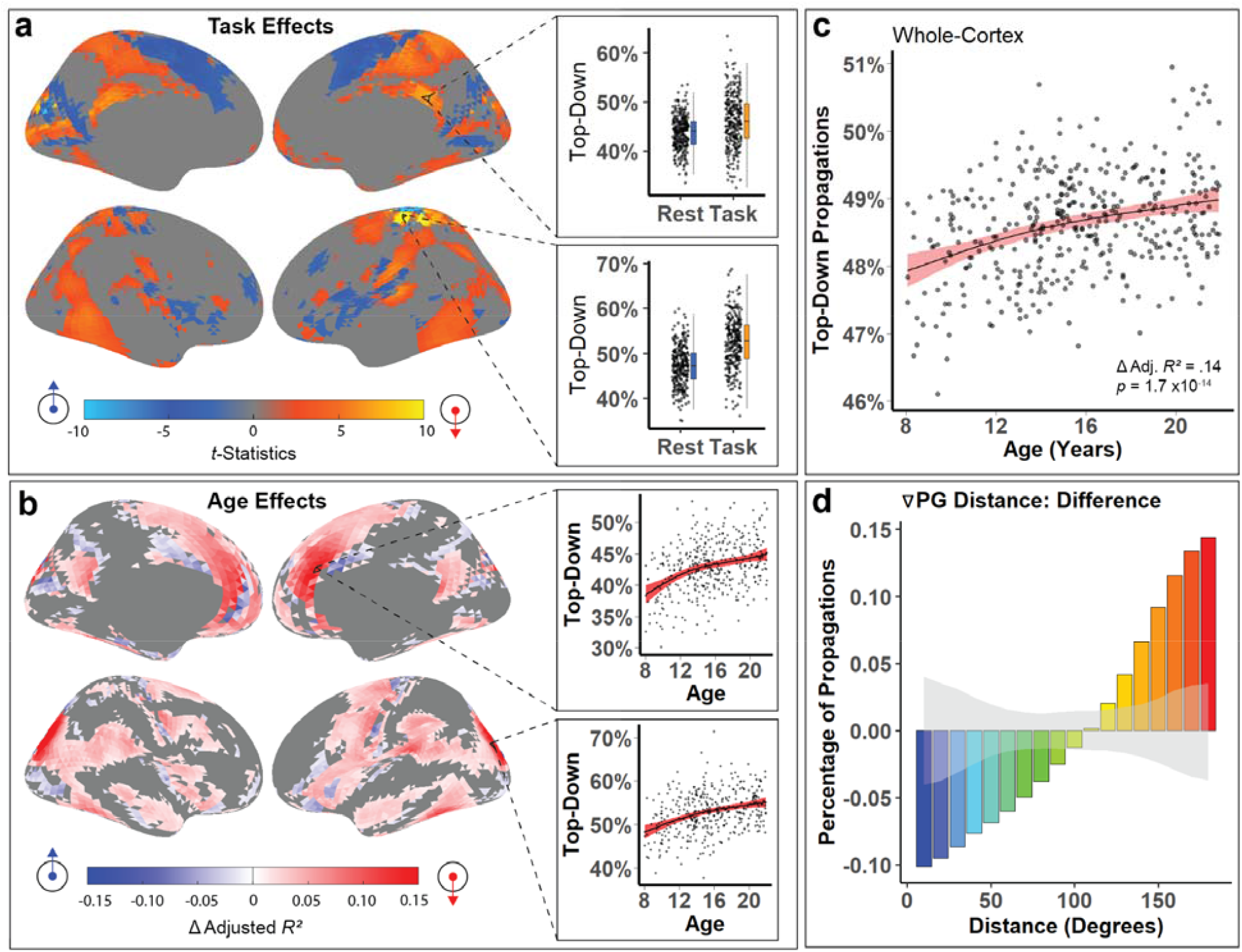
The prevalence of top-down propagations is impacted by task demands and develop with age. **a)** Compared to rest, the demands of a cognitive control task elicit a shift in the proportion of propagations that are top-down (*P*_*FDR*_ < 0.05, more top-down under task demands in orange). **b)** Top-down propagations become more prominent with age in youth, particularly in attention systems (*P*_*FDR*_ < 0.05, more top-down with age in red). **c)** When averaged across the cortex, top-down propagations increase with age (smooth term effective degrees of freedom = 1.89). **d)** Whole-cortex directional distributions mature such that after adolescence, a greater percentage of propagations are top-down. This difference extends above and beyond distribution differences observed in 1,000 equally sized, randomly selected subgroups of participants (gray band = 95% confidence interval on bootstrap resamples).

Next, we evaluated whether the prevalence of top-down propagations evolved with age in youth. Analyses using generalized additive models that capture both linear and nonlinear effects revealed widespread increases in the proportion of top-down propagations observed with age across the cortex (Δ Adjusted *R*^*2*^ = 0.01 - 0.19, *p*_fdr_ <0.05, **Figure 3b**). These effects were particularly prominent in the dorsal and ventral visual streams, as well as the medial and lateral premotor pathways. Surprisingly, age effects extended continuously beyond the canonical premotor pathway into inferio-medial prefrontal cortex. These results suggest that maturation of internally-oriented default-mode regions may be spatially overlapping with maturation of the internally-driven medial premotor pathway^22^. These observations were part of a broader pattern of increases in the proportion of top-down propagations across the cortex (Δ Adjusted *R*^*2*^ = 0.14, *p* = 1.7×10^−14^, **Figure 3c**).

We next sought to determine how development alters the full distribution of propagation directions rather than simply evaluating the change in proportion of top-down or bottom-up flow. To do so, we calculated the difference in the average angular distribution of propagations for the youngest (*n* = 127, mean age = 11.49, *SD* = 1.70 years) and oldest (*n* = 132, mean age = 19.76, *SD* = 1.39 years) tertile of the data (**Figure 3d**). We then evaluated the significance of this difference of distributions by comparing the true difference to a null distribution created from random tertile splits (**Figure 3d**, gray band). We found that the angular distributions shift monotonically towards top-down propagations with age: maximally top-down propagations increased with age the most, whereas maximally bottom-up propagations showed the largest declines with age.

Finally, we conducted sensitivity and specificity analyses to confirm our findings. Notably, the spatial distribution of the principal gradient is collinear with the distribution of functional networks^10^, and the age effects we report occur over the same age range as developmental functional network segregation^6^. To ensure that our developmental results were not attributable to previously reported functional network segregation, we quantified network segregation in all participants. While controlling for network segregation, increases in top-down propagations over development remained prominent (Δ Adjusted *R*^*2*^ = 0.14, *p* = 1.2×10^−14^, **Figure S4**) and exhibited a stronger age-effect size than network segregation itself (Δ Adjusted *R*^*2*^ = 0.05, *p* = 3.1×10^−5^). Finally, to verify that age effects were not attributable to scanning-site differences, we performed ComBat harmonization and repeated the above analyses. Developmental effects remained prominent when accounting for site differences (Δ Adjusted *R*^*2*^ = 0.12, *p* = 2.0×10^−12^, **Figure S5**). Together, these sensitivity and specificity analyses confirmed that our findings were not attributable to previously documented properties of functional neurodevelopment or scanner differences.

Several limitations should be noted. First, the temporal resolution of fMRI restricted our analyses to infraslow frequencies. Although animal studies continue to corroborate the importance of infraslow fluctuations in brain-wide neuronal synchronization and behavioral arousal^23,24^, future studies may reveal similar^15,16^, independent^24^, and inverted^25^ propagation patterns in different frequency domains. Second, the cost function of optical flow is agnostic to the positivity of the propagating signal: propagating *decreases* in BOLD signal are also captured by the resulting vector fields. Because hierarchically propagating infraslow activations and deactivations can facilitate or suppress faster rhythms^15,16^, explicitly delineating activations from deactivations is an important step for future work. Third, respiratory^16,26^ and vascular^18,27^ processes are known to contribute to BOLD signal propagations at similar frequencies. Further analyses of concurrently acquired physiological data^15,16,28^ may serve to disentangle their contributions to observed propagations. Nonetheless, existing evidence suggests that traveling local field potentials may underlie the propagations observed in this study^29-33^. Finally, motion-related signal artifact is likely to have a substantial impact on functional propagations. Consequently, we erred on the side of being extremely stringent in quality assurance – using only low motion data and statistically controlling for residual motion artifact in all analyses.

These limitations notwithstanding, we developed an approach to quantify how propagations align with the cortical hierarchy. This revealed that activity preferentially flows up and down the cortical hierarchy. Our observation that top-down propagations increase in response to top-down task demands suggests that such propagations are to some degree state-dependent. This observation coheres with initial evidence from other studies^12,14^, and further suggests that top-down processing may rely upon hierarchical cortical propagations. Finally, we found that top-down propagations become more prominent with age in youth. Our findings suggest that the directionality of propagating cortical activity may be broadly relevant for studies of hierarchical cortical organization and neurodevelopment, with potentially important implications for our understanding of psychopathology and the design of neuromodulatory interventions.

## Methods

### Sample

To evaluate the maturation of cortical propagations, we used high-quality resting-state and task-fMRI data from the Human Connectome Project-Development 2.0 Release (HCP-D, *n* = 652, mean age = 14.4, *SD* = 4.1 years). Participants were scanned at four sites on 3 Tesla Siemens Prisma platforms. Structural scans consisted of high-resolution MPRAGE T1w images (0.8 mm^3^, TR/TI=2,500,1000 ms, TE = 1.8/3.6/5.4/7.2 ms, flip angle = 8°) and a variable-flip-angle turbo-spin-echo T2w sequence (0.8 mm^3^, TR/TI=3,200,564 ms, turbo factor = 314). Additionally, each subject underwent 26 minutes of resting-state scans across 4 runs, and 8 minutes of task-fMRI across 2 runs for our task of interest^21^. Multiband acceleration factors afforded sub-second temporal resolution for all functional images (2.0 mm^3^, TR/TE = 800/37 ms, flip angle = 52°).

### Image processing

All images were processed with an updated version of the Human Connectome Project MRI pipeline^34,35^. Specifically, all structural images underwent gradient distortion correction, bias field correction, boundary-based registration, and normalization. Functional images underwent gradient distortion correction, re-alignment, EPI distortion correction, boundary-based registration, and normalization prior to being projected to the cortical surface and smoothed with a 2mm FWHM gaussian kernel. Next, functional images were demeaned and de-trended using nuisance regressors. Finally, functional images were band-pass filtered between 0.008 and 0.09 Hz with a 2^nd^ order Butterworth filter. Framewise displacement was calculated after accounting for the influence of respiratory signal on framewise image realignment. Noteworthy changes from the HCP pipeline included usage of Advanced Normalization Tools (ANTs) for denoising, bias field correction, and diffeomorphic symmetric image normalization, which was selected due to consistently higher registration performance over previous methods^36^. Finally, all images were downsampled to *fsaverage4* with connectome workbench for computational feasibility.

### Quality assurance

In order to be included in analyses, participants needed to have at least 600 TRs surviving three quality-control thresholds. First frames were excluded if head motion exceeded 0.2 mm framewise displacement for that frame. Second, frames were excluded if they contained DVARS values that were > 3 standard deviations above the mean. Third, because we were interested in propagations across TRs rather than patterns within single, low-motion TRs, we excluded otherwise low-motion segments that were interrupted by moderate to high-motion frames. Specifically, if TRs that met the first two criteria were not part of a broader sequence of at least 10 consecutive low-motion TRs, these TRs were discarded. 388 participants (mean age = 15.6, *SD* = 3.7 years) met the > 600 TR requirement after the aforementioned quality assurance procedures.

### Cognitive control task

For task-fMRI, we selected the Carit task *a priori* because it requires top-down cognitive control. The Carit task is a modified Go/No-Go task, where participants are instructed to make repeated button-presses in response to rapid, consistent stimuli, which are periodically interrupted. At the time of this interruption, the participant is to withhold a button press, probing their ability to suppress their button-pressing response. Because fewer scans were allocated to this task within HCP-D, we relaxed the minimum TR requirement to 300 TRs for task analyses only. As we compared propagations between task and resting conditions on a within-subject basis, only participants who passed both resting-state quality control (600 remaining TRs) and task QC (300 TRs) were included for these analyses.

### Optical flow

Optical flow is a computer vision technique used to estimate the motion of signal intensity between successive images^37^. Like image registration, this procedure optimizes the deformation field that best explains the spatial discrepancy of signal intensity between two images. Recently, the optical flow algorithm was adapted to efficiently estimate biological motion on spherical surfaces^20^. We leveraged this advance to quantify the movement of the BOLD signal directly on each participant’s cortex following spherical registration. As 2-dimensional “patch” projections of the cortex incur large discontinuities between spatially adjacent cortices, use of the spherical implementation of optical flow allowed us to efficiently analyze propagations across the cortex.

### Defining hierarchical ascent and descent

In order to estimate directions of hierarchical ascent and descent, we extracted the gradient vector field (∇) of an established map that defines the principal gradient (PG)^13^ of the cortical hierarchy (∇PG). This approach is analogous to that taken in Tian *et al*. (2020)^38^, but extracted across the cortical mantle rather than in subcortical volumetric space.The resultant vector field, describing hierarchical ascent at each face on the cortical mesh, was subsequently used as a common set of reference directions for each participant’s optical flow data.

### Quantification of angular distances

In order to evaluate directional alignment between optical flow vectors and hierarchical vectors, we evaluated their angular similarities in degrees. Magnitude measurements were discarded from optical flow and ∇PG; only directional information was reported. Our primary metric of interest was the angle (in degrees) between hierarchical vectors and optical flow vectors. To derive these angles, the 3-dimensional cartesian (x,y,z) vectors describing both vector fields were converted to a spherical coordinate system (azimuth, elevation, rho) via *cart2sphvec* in MATLAB. Because the signal travels across the surface of the sphere rather than into or away from it, this conversion obviates the third coordinate (rho). Consequently, we retained azimuth and elevation only for each hierarchical and optical flow vector, which describe directionality on a 2-D tangent-plane at each cortical face **(Figure 1c)**. From this point, the angular distance was computed as the difference in directional orientation in degrees between ∇PG and optical flow, with 0 degrees indicating perfect alignment and 180 degrees indicating the maximum possible difference.

### Assessment of alignment between ∇ PG and null models

In order to test whether hierarchical ascent and descent were both directional modes in the distribution of optical flow vectors, we employed Hartigan’s dip test. Specifically, we used the dip statistic to quantify the deviance of angular distributions from a unimodal distribution: a higher dip stat indicates that a distribution is more likely multimodal than unimodal. Subsequently, we compared this measure to dip statistics derived from spatial and temporal null models.

For spatial null models, optical flow angular distances were calculated relative to a spatially “permuted” ∇PG. By rotating or “spinning” the entire ∇PG continuously in space, local spatial properties of the original map are conserved^39^. Consequently, this procedure yields a more realistic and conservative spatial null model than random permutations where the spatial covariance structure is lost. We performed 1,000 permutations, and 1,000 corresponding null dip statistics were obtained for each participant. Finally, to extract a metric comparable across participants, we recorded the number of standard deviations between the true observed dip statistic and the mean of the 1,000 permutations.

For temporal null models, optical flow itself was re-calculated on temporally permuted data. Specifically, the temporal sequence of fMRI volumes surviving QC was shuffled iteratively for each participant. Because fitting optical flow to a pair of frames is computationally intensive (equivalent to a co-registration), we were limited to 100 temporal permutations per subject (613,000-1,883,000 optical flow decompositions per subject). This process yielded 100 sets of optical flow vectors for each participant’s shuffled data. These null sets of vectors were then subjected to the same angular distance calculation (relative to ∇PG), and 100 null dip statistics were subsequently obtained from these distributions. As for the spatial permutation tests, we compared true vs. permuted dip statistics as a single participant-level standard deviation.

### Analysis of the impact of task demands

To test our hypotheses regarding shifts in top-down propagation prominence with task, we quantified the proportion of propagations that descended the cortical hierarchy. To do so, we calculated the proportion of optical flow vectors that indicated descent in any capacity (i.e., greater than 90 degrees from ∇PG) versus optical flow vectors that indicated hierarchical ascent (i.e., less than 90 degrees from ∇PG). This procedure provided a measure of the prevalence of top-down propagations at each cortical face for each participant.

We then compared the proportion of top-down propagations during rest and under the cognitive control demands of the Carit Task. Specifically, we conducted a paired *t*-test on the proportion of top-down propagations at each cortical face. This procedure provided a *t*-statistic quantifying the degree to which faces exhibited more top-down propagations during the task than during rest. Multiple comparisons were controlled by the false-discovery-rate (FDR: *q* < 0.05); only statistics that remained significant after correction for multiple comparisons were retained and reported.

### Analysis of developmental effects

Developmental effects were estimated using generalized additive models^40^ (GAMs) with penalized splines in R (Version 3.6.3) using the *mgcv* package. Non-linearity was penalized to avoid over-fitting, and fitting was optimized with restricted maximum likelihood (REML)^41^. Participant sex, in-scanner head motion, and the number of frames passing quality assurance were included as covariates within each GAM. Four knots were specified as the maximum flexibility afforded to age splines in all models. To quantify the effect sizes of each age spline, we calculated the change in adjusted *R*^2^ (Δ*R*^2^_*adj*._) between the full model and a nested model that did not include an effect of age. Statistical significance was assessed using analysis of variance (ANOVA) to compare the full and nested models^42^. As above, multiple comparisons were controlled for with the false-discovery-rate (*q* < 0.05). Finally, because Δ*R*^2^_*adj*._ describes effect size but not direction (i.e., increasing or decreasing top-down propagations with age), as in prior work^6^, we extracted and applied the sign of the age coefficient from an equivalent linear model.

To quantify developmental differences in the full distributions of angular distances between optical flow vectors and ∇PG, we compared the oldest and youngest tertiles of all participants. Specifically, we reduced each participant’s angular distribution to 18 bins, with each bin comprising a 10-degree span from 0-180 degrees from ∇PG. Each bin represents the percentage of optical flow vectors that fell within a 10-degree window of angular distances from ∇PG (0-10 degrees, 10-20 degrees, etc.) Across participants within each tertile split, the average of these percentages represents the average percentage of total propagations each 10-degree bin encompasses for each age tertile. To observe age-dependent differences, we subtracted the resultant value of each bin in the younger tertile from the resultant values in the older tertile. This approach provided a description of the difference in angular distributions between older and younger participants. However, that difference measure does not provide a statistical test of whether the difference is significant. To evaluate the statistical significance of age effects, we performed a bootstrap procedure, where tertile splits were determined randomly. We repeated the difference-of-distributions procedure described above for 1,000 random tertile splits, producing 1,000 random differences of distributions. Finally, we extracted the 95% confidence interval from these 1,000 distribution differences to obtain an estimate of distribution differences that could be expected by chance alone. Observed differences exceeding this confidence interval were interpreted as true group differences, exceeding those expected by selecting two groups of the same size when the age distribution was random.

### Sensitivity and specificity analyses

We used sensitivity analyses to confirm that our results were not due to confounding factors. First, to ensure that hierarchical development of cortical propagations is not explained by hierarchical development of cortico-functional networks, we repeated our analyses while controlling for developmental network segregation. To do so, we constructed a 17-network group-consensus atlas for the participants in our study with spatially regularized non-negative matrix factorization. Next, we calculated network segregation as prior^6^: the mean between-network coupling of a network with all other networks. We included this value as a model covariate in sensitivity analyses. Based on prior work, we identified which of the delineated networks are those most likely to exhibit developmental segregation. Previously, we have detailed that the functional networks undergoing the most dramatic developmental segregation are those lying at the top of the cortical hierarchy^6^, and other publications have similarly suggested that default-mode networks undergo developmental segregation^43,44^. Accordingly, we evaluated each network for its hierarchical position and overlap with canonical functional networks, and selected the single network fulfilling both *a priori* criteria (high in hierarchy and overlapping with the canonical default mode). Segregation of this network (higher-order default-mode) comprised the metric of interest for our first sensitivity analysis.

Finally, to ensure that the association between top-down propagations and age were not attributable to site effects, we harmonized top-down propagations across sites with ComBat^45,46^. This provided a site-harmonized measure of the proportion of top-down propagations exhibited by each participant, which we then tested in the same GAM framework.

### Code availability

All analysis code and a step-by-step replication guide is available at https://github.com/PennLINC/PWs.

## Supplemental Figures

**Figure S1:**
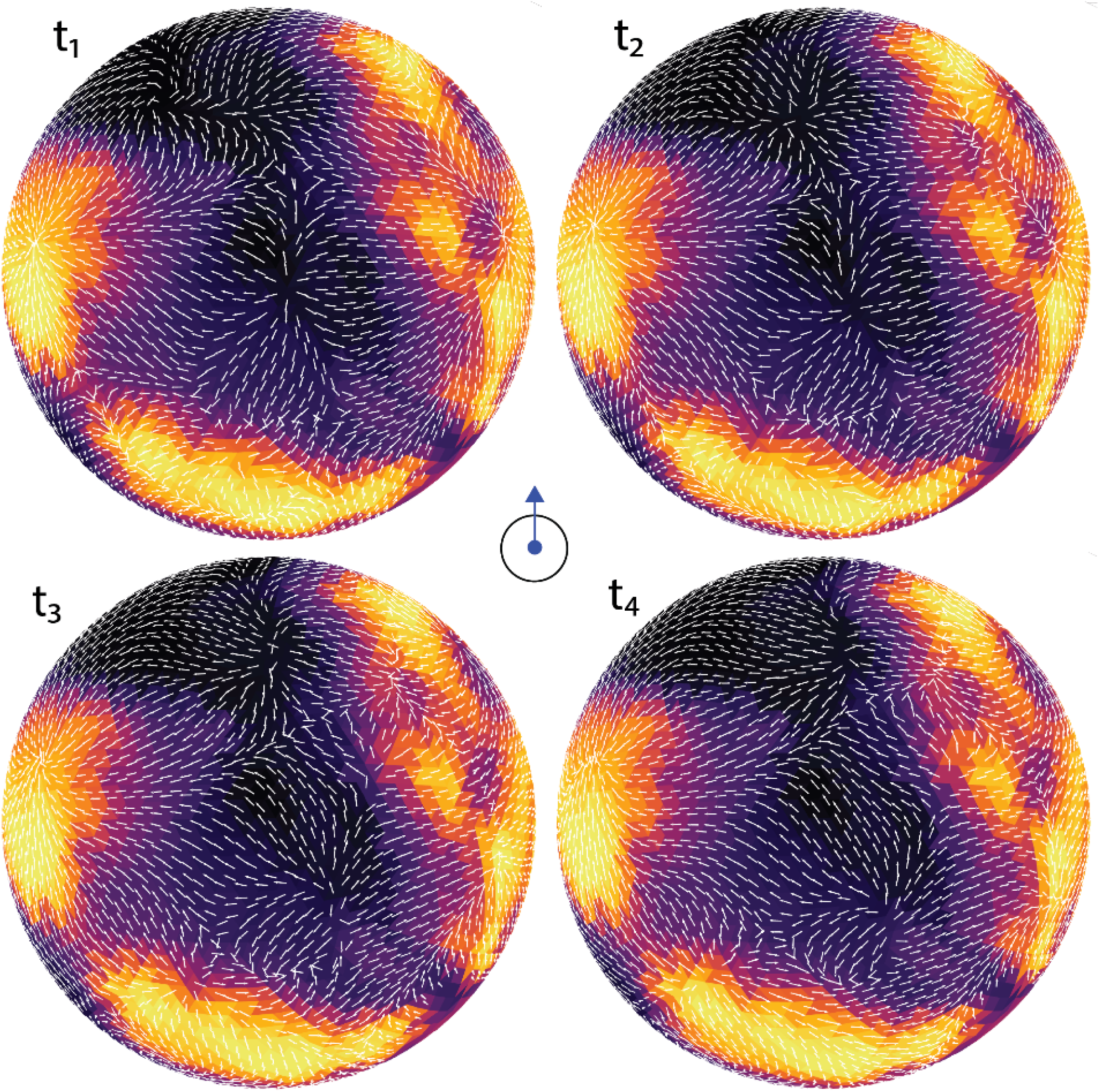
Whole-hemisphere visualization of the bottom-up propagation sequence depicted in **Figure 1e** in the main text.

**Figure S2:**
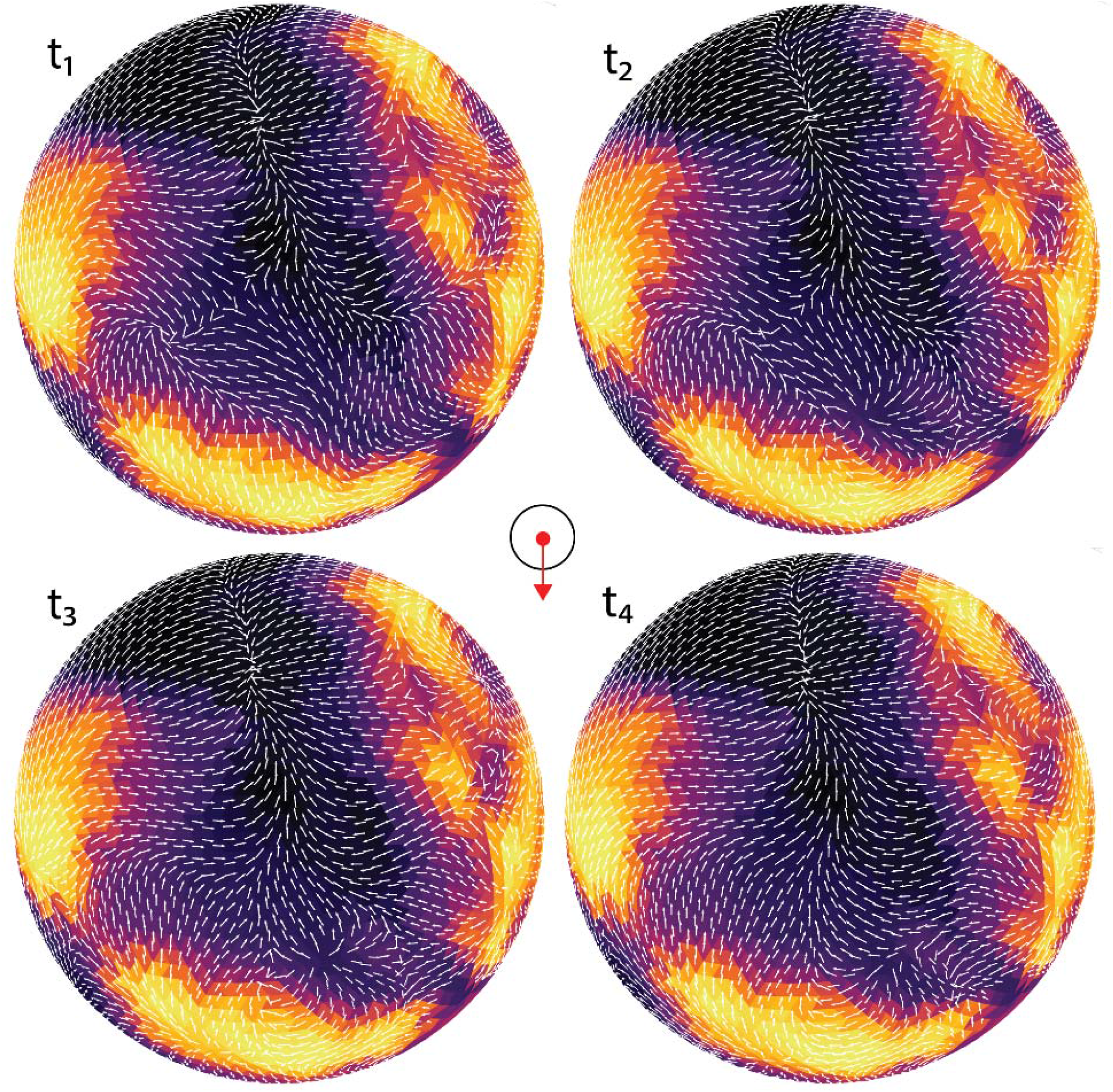
Whole-hemisphere visualization of the top-down propagation sequence depicted in **Figure 1e** in the main text.

**Figure S3:**
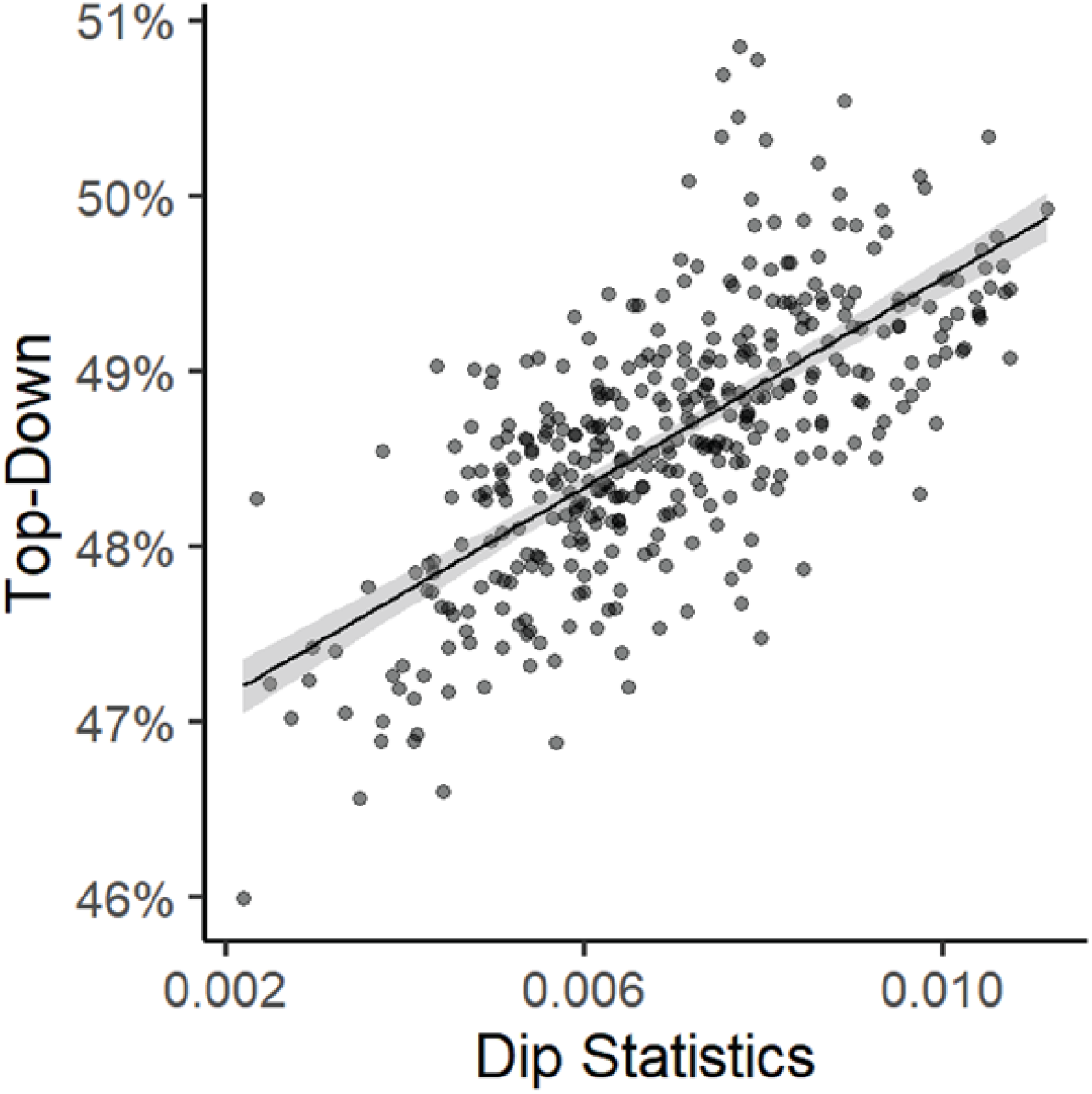
Across participants, top-down propagations are strongly associated with non-unimodality (e.g., the dip statistic) of directional distributions of optical flow vectors (*r* = .70, *p* < 0.01×10^−14^). The 95% confidence interval of this linear relationship is indicated by the shaded area.

**Figure S4:**
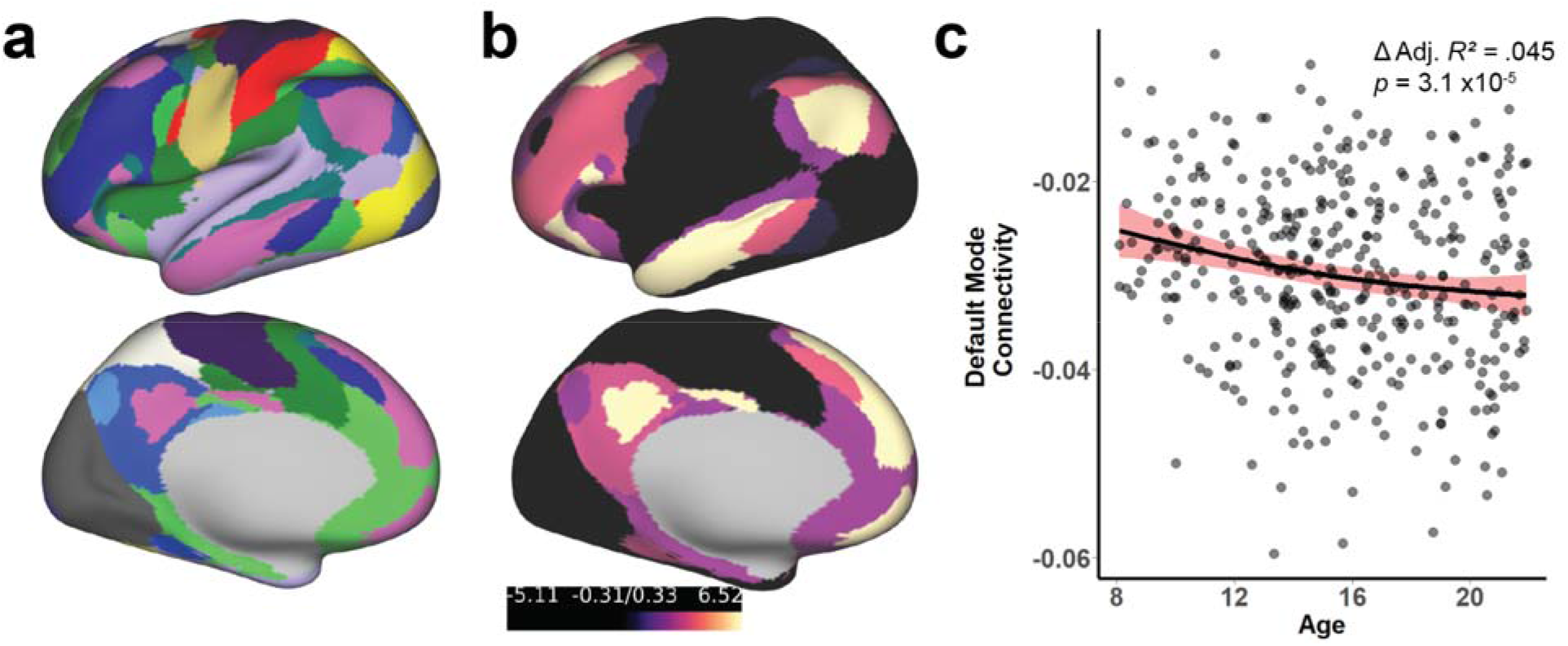
Hierarchical functional network segregation is also related to age. **a)** Group consensus functional network template derived for HCP-D with regularized non-negative matrix factorization. Functional network membership is indicated by color. **b)** Each network ranked by the average value of the principal gradient with that network. Hierarchical positioning indicated by brighter coloration. Only one network (magenta in panel **a**, white in panel **b**) fulfilled both *a priori* criteria of default-mode overlap and high hierarchical positioning. **c)** Consistent with our prior work, this network exhibited segregation from all other networks over development, even after controlling for the proportionate increases in top-down propagations observed with age (**Figure 3;** smooth term effective degrees of freedom = 1.71). Shaded area indicates 95% confidence interval on the smooth term estimated for age.

**Figure S5:**
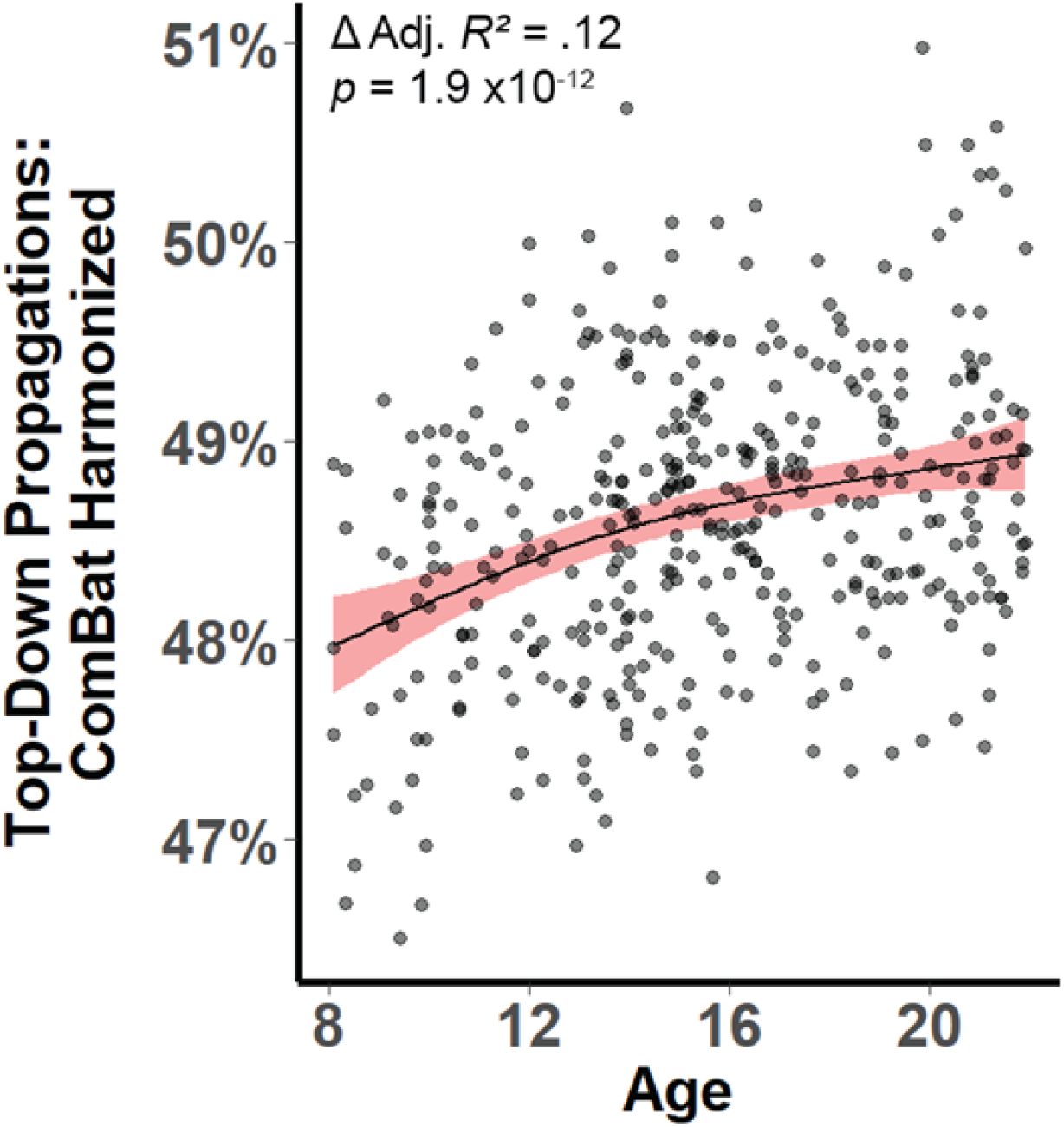
Observed top-down propagation development is not driven by site effects. Top-down propagation maturation is substantial after controlling for site with ComBat harmonization (Δ Adjusted R^2^ = 0.12, *p* = 2.0×10^−12^, smooth term effective degrees of freedom = 1.87). Shading indicates 95% confidence interval for estimation of the age spline.

